# The neuro-ocular costs of texting during driving

**DOI:** 10.1101/2025.08.15.670515

**Authors:** Tzvetan Popov, Nan Li, Veronika Gambin, Kristina Keller, Samuel Wehrli, Stefan Lakämper

**Affiliations:** Methods of Plasticity Research, Department of Psychology, University of Zurich, Zurich, Switzerland; Department of Psychology, University of Konstanz, Konstanz, Germany; Division of Traffic Medicine, Institute for Forensic Medicine, University of Zurich, Zurich, Switzerland; School of Life Sciences and Facility Management, Zurich University of Applied Sciences, Research Group Biosensor Analysis and Digital Health, Zurich, Switzerland

## Abstract

Texting while driving is among the most dangerous forms of distraction, yet the neurophysiological mechanisms linking cognitive load, gaze control, and driving behavior remain poorly understood. Here, EEG, eye tracking, head kinematics, and driving performance were recorded in a naturalistic driving task while participants solved arithmetic problems presented on a dashboard-mounted display under low and high working-memory (WM) load, mimicking real-world interaction with dashboard media. High WM load resulted in slower reaction times, reduced accuracy, prolonged gaze toward the stimulus, increased corrective head rotations, and compensatory reductions in driving speed. Time-frequency analysis revealed robust alpha power suppression (∼8-13 Hz) over occipito-parietal regions scaling with WM load and closely paralleling gaze behavior. Source reconstruction further identified a transient, stimulus-locked increase in oscillatory power at ∼15 Hz involving the ipsilateral cerebellum and posterior parietal preceding the head rotation onset. These findings illuminate the time course of a neuro-ocular action circuit involving cerebellum and cortex as candidate neurophysiological precursor of upcoming behavioral costs as a result of driver distraction. Bridging cognitive neuroscience and applied traffic research, the present results provide an ecologically valid framework for studying action-attention coupling in real-world behavior.

## Introduction

Texting while driving remains one of the most dangerous forms of driver distraction. Yet, interacting with dashboard media is an integral part of the contemporary driving experience. Actions such as adjusting the air conditioner, tapping a smartphone in response to a Google Maps prompt, or briefly engaging with social media each represent varying levels of distraction that require careful contextual assessment. Among these, texting while driving constitutes a particularly multifaceted distraction, simultaneously taxing the driver’s visual, manual, and cognitive resources. However, exactly how the visual and cognitive components interact to impair driving performance remains unclear. By the time a texting driver visibly drifts out of their lane or narrowly avoids a collision, substantial distraction-induced impairment has already occurred. Thus, identifying early neuro-cognitive and ocular indicators of attentional lapses-before observable driving errors occur-is a critical yet elusive research objective^1,2^.

Driver distraction involves several cognitive processes: prioritizing tasks (e.g., focusing on the road versus attending to the dashboard), allocating attention and encoding stimuli (e.g., reading a text message), and maintaining encoded information to guide subsequent actions (e.g., immediately responding or delaying the response). This sequence closely parallels classic models of attention and working memory studied extensively in controlled cognitive neuroscience laboratories. Specifically, prior research emphasizes the role of fluctuations in alpha oscillatory power for allocating spatial attention, maintaining working memory and preventing incoming distractors [e.g. ^3–9^]. Attention deployment typically coincides with alpha power reductions over the occipito-parietal cortex, and encoding or maintaining higher working-memory loads similarly correlates with greater alpha suppression^10–13^. Recent report highlights that these power modulations function as a cortical mechanism to suppress distraction as perceptual load increases^14^, a phenomenon that is highly relevant to many real-world situations, among them eminently, driving [e.g. Figure 1 in^3^]. While the theoretical interpretation of these findings remains debated^5,15–18^, their reproducibility^19^ provides a clear empirical prediction: elevated attentional demand and working-memory load in the context of driver distraction should be associated with decreased alpha power. In this case, given the predominant view on the functional role of alpha oscillations, a plausible interpretation is that the decrease in alpha power reflects a release of inhibition, thereby facilitating the activation of cortical circuits that prioritize relevant and filter out irrelevant information within the conceptual framework of top-down cognitive control. Not surprisingly, a similar interpretation has already been proposed in the context of driving^20–22^. While intuitively appealing, this account remains difficult to substantiate, as it implicitly relies on a homuncular explanation for why and how such a complex cognitive construct is actually implemented.

**Figure 1:**
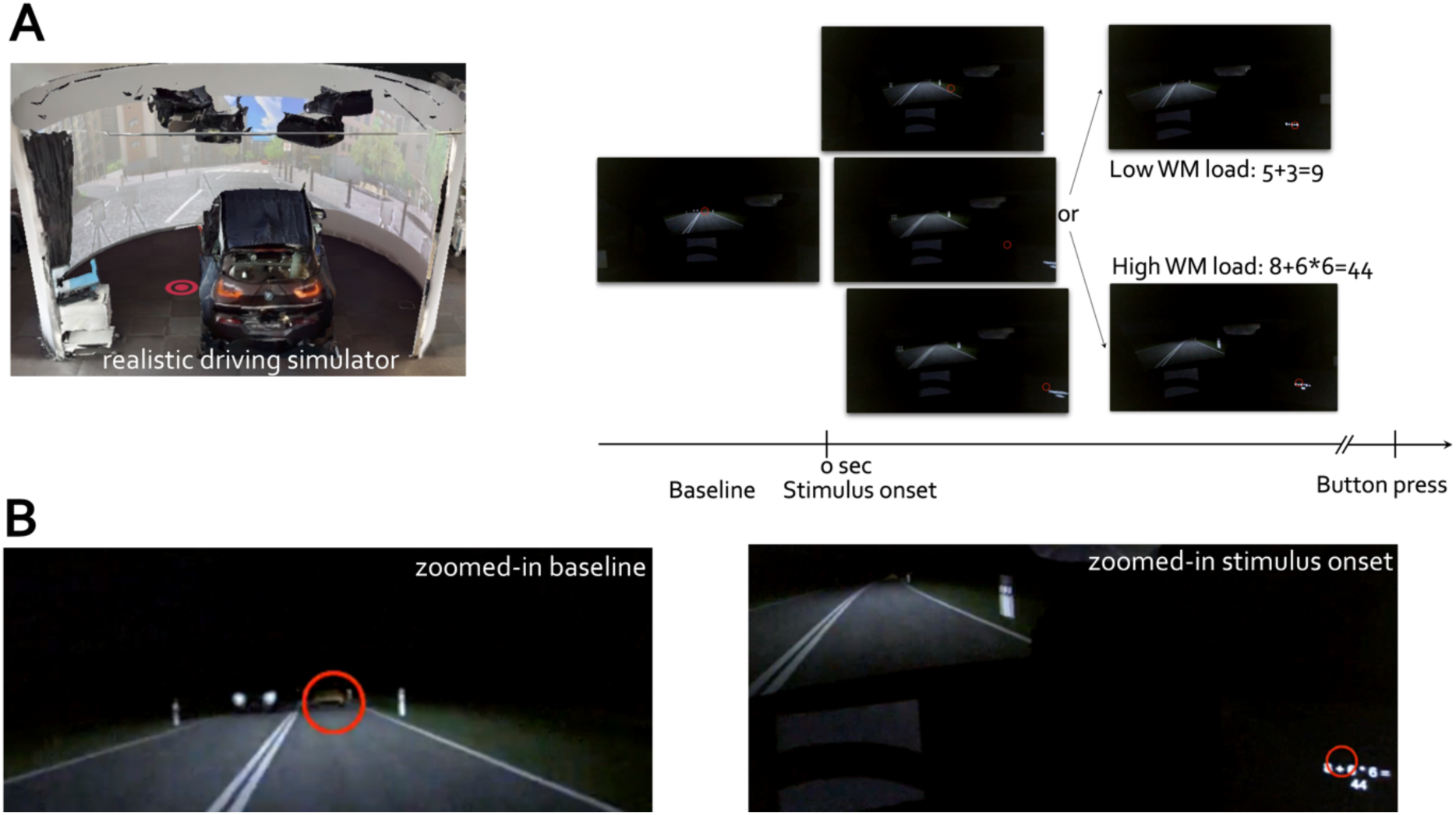
Dual-task paradigm combining simulated driving with an arithmetic working memory (WM) task, illustrating performance decrements under high WM load. **A-** Illustration of the realistic driving simulator. Schematic timeline of a trial, showing an initial baseline period followed by stimulus onset and presentation of an arithmetic problem of either low (e.g., 5 + 3 = 9) or high WM load (e.g., 8 + 6 × 6 = 44). Participants had to indicate via button press whether the presented solution was correct (one space key press) or incorrect (double space key press) while navigating a simulated driving environment. Red circles indicate the participant’s gaze position (point of view, 1dva) in the driving scenes. **B-** zoomed-in views of the gaze positions for the baseline (orange) and stimulus (green) periods, highlighting the precise visual focus of participants during each task phase.

A complementary perspective arises from considering the primacy of action-specifically gaze behavior-in enabling perception and directing attention toward objects in the environment^23–25^. This perspective follows from the fact that the default mode of the human eye is movement. Humans produce about 150,000 saccades per day^26^, which amounts to approximately 1.5 hours of daily oculomotor activity dedicated solely to these rapid gaze shifts. In addition, the eyes are never truly at rest; they continuously generate microsaccades, drifts, and tremor during fixation, ensuring that the visual system remains dynamically engaged throughout nearly the entire waking day^27,28^ and even continued during sleep. This constant interplay between saccadic and fixational eye movements underscores that vision is inherently active, with gaze behavior forming the primary interface through which perception and attention are directed toward the external world. And yet, this fact is rarely acknowledged in laboratory experiments^23^. Instead, a static interval-during which participants are typically asked to fixate on a central cross-is treated as „at rest“-baseline against which experimental data are compared. When time-frequency representations of power are analyzed relative to this baseline, they almost invariably reveal a decrease in alpha power from the pre-stimulus interval, regardless of the context or cognitive construct under investigation. This prevailing approach relies on the very assumption that a) static fixation exists, b) can be experimentally imposed on the observer, and therefore c) constitutes a valid contrast (i.e. free from eye movements and cortical control thereof) for evaluating experimental intervals. However, if one accepts the premise that the default mode of the eye is movement, then a mechanism must be postulated that can temporarily suppress this intrinsic oculomotor activity, reducing the degrees of freedom of eye movements, and thereby aiding a temporal manifestation of “fixation”, and enhancing visual acuity. This requirement naturally points to alpha oscillations as a plausible neural mechanism for regulating this default oculomotor mode^17,23,29–32^, with the 100 ms long duty cycle of alpha oscillations corresponding to the temporal window required for initiating an eye movement mediated by the superior colliculus and cortex^33^. Rather than reflecting purely cognitive inhibition, alpha activity may serve to transiently stabilize gaze, suppress exploratory eye movements, and thereby create the artificial state of enforced fixation that is used as the conventional baseline in experimental paradigms. Conversely, an increase in oculomotor exploratory behavior is associated with a depression of alpha power, a conjecture that is receiving increasing empirical support across diverse domains, including visual and auditory spatial attention^34,35^, picture and face viewing^23^, and episodic memory encoding^36,37^. The commonly observed pattern is that decreases in alpha power are inversely related to oculomotor exploration-the greater the exploration, the stronger the alpha suppression. This perspective reframes alpha oscillations not as an abstract marker and somewhat passive manifestation of “top-down control”^38–40^, but as an active neural mechanism for regulating the default oculomotor state, thereby not only eliminating the need for a homuncular agent but also allowing for grounding attentional control in measurable sensorimotor dynamics.

Driver distraction frequently initiates through changes in oculomotor behavior. EEG has long been recognized as a powerful tool for investigating cognitive load and attentional states during driving, and recent reviews have highlighted its potential for detecting driver distraction and supporting intelligent assistance systems^1,2^. However, most existing studies have relied on simplified laboratory paradigms, often using above-mentioned static fixation and isolated secondary tasks that fail to capture the complexity of real-world driving. Moreover, while hybrid approaches combining EEG with eye tracking have been recommended, few studies have explicitly linked neural activity-particularly alpha oscillations- to naturalistic gaze and head movements. This has reinforced a predominantly cognitive interpretation of alpha power as a marker of top-down inhibition, while largely overlooking its emerging role in sensorimotor control and visuospatial action.

To test the hypothesis that distraction-induced attentional shifts-manifesting as spatially specific gaze variability-are accompanied by concurrent decreases in alpha power in a naturalistic setting, the present proof-of principle study combined recording EEG, eye tracking and head kinematics with a real world-related behavioral performance task during naturalistic driving in a highly immersive driving simulator. Participants performed a dual-load working-memory task while simultaneously driving in a state-of-the-art realistic driving simulator (https://my.matterport.com/show/?m=ntULEgy28cU). EEG, eye-tracking, and head-movement data were collected concurrently, along with critical driving parameters such as lateral position, steering-wheel deviation, and accelerator-pedal pressure. Stimuli were presented centrally on a dashboard-mounted screen in two distinct working-memory load conditions: a low-load condition involving simple arithmetic (addition and subtraction), and a high-load condition involving more complex arithmetic requiring adherence to the order of operations. Participants indicated via button press whether the displayed solution was correct or incorrect, thereby creating a realistic yet minimally distracting scenario analogous to responding to prompts such as Google Maps queries.

The tested predictions were as follows: at the behavioral level, reaction times were expected to increase and accuracy to decrease under higher working-memory load. At the neural level, alpha power was predicted to exhibit a stronger decrease prior to button presses in high-load compared to low-load conditions. Furthermore, greater gaze variability was anticipated under higher load, paralleling the expected suppression of alpha power.

## Results

Point-of-view (POV) illustration of the task design is depicted in Figure 1. Participants were instructed to follow a car along a single-lane highway while maintaining fixation on the road ahead (indicated by the red circle representing eye-tracking gaze position). In the context of the present study, this interval was considered the baseline condition. At random intervals, a stimulus (math problem solution) was presented on a screen positioned centrally on the right-hand side of the driver, under two conditions requiring either low or high working memory (WM) load. The presentation of the stimulus elicited a gaze shift away from the road toward the stimulus location, as illustrated by the shift in the red circle in Figure 1. Participants were instructed to judge whether the solution was correct by pressing a button once if it was correct or twice if it was incorrect (Figure 1A, right). The solutions either required low WM load (e.g., *5 + 3 = 9*) or high WM load (e.g., *8 + 6 × 6 = 44*). A total of 200 stimuli (100 per condition) were presented in random order over the course of the driving session, which lasted approximately 45 minutes.

The variation in WM load affected both reaction time and accuracy (Figure S1, Supplementary material). Participants responded faster in the low-load condition (M = 3.2 s, SD = 1.2) compared to the high-load condition (M = 4.5 s, SD = 1.5). This difference was statistically significant, as confirmed by a two-sided t-test, *t*(42) = −11.2, *p* = 3.49 × 10⁻¹⁴, SD = 0.8, 95% CI [−1.6, −1.1]. Similarly, accuracy was higher in the low-load condition (M = 97.7%, SD = 2.1) compared to the high-load condition (M = 95.8%, SD = 3.2)-*t*(42) = 4.5, *p* = 4.56 × 10⁻⁵, SD = 2.8, 95% CI [1.1, 2.8]. Figure 2 summarizes the recorded time series of biological and driving data. Stimulus onset elicited a transient event-related potential (ERP; Figure 2A) over occipital-parietal electrodes (P7, P3, P4, P8, POz, O1, O2, M1, M2), peaking approximately at 360 ms after stimulus onset. This was followed by a sustained potential lasting several seconds, which was significantly stronger for the high WM load condition compared to the low WM load condition, as confirmed by a cluster-based permutation test (*p* < 0.05; Figure 2A and Figure S2A in the Supplemental material). Gaze velocity increased immediately after stimulus onset, peaking as early as 200 ms (Figure 2B). This was followed by a sustained increase in gaze velocity for the high WM load condition compared to the low WM load condition, closely paralleling the sustained activity observed in the evoked cortical response. This effect was confirmed by a cluster-based permutation test (*p* < 0.05; Figure 2B). Head rotation velocity (around the vertical Y-axis measuring yaw) toward the stimulus on the dashboard peaked approximately 1 s after stimulus onset, followed by a rotation in the opposite direction-back toward the road-around 2 s. Compared to the low WM load condition, the angular velocity of head rotation back toward the road was weaker in the high WM load condition, indicating prolonged maintenance of head orientation toward the stimulus rather than the road. Additionally, repeated small head rotations persisted later in the trial under high WM load (Figure 2 C, cluster-based permutation test p <0.05).

**Figure 2:**
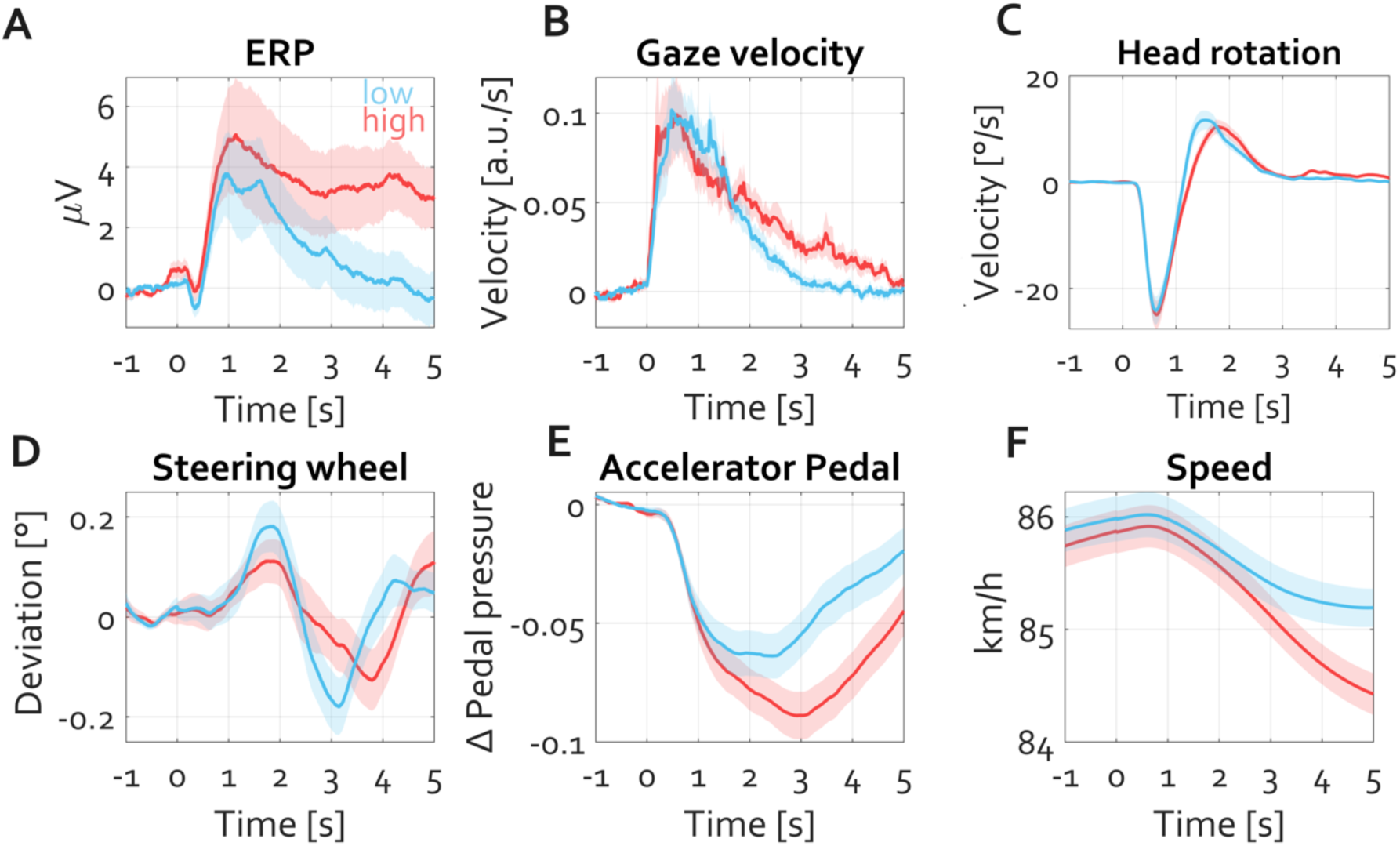
Neural, ocular, head, and driving-related responses as a function of working-memory (WM) load. **A**- Event-related potentials (ERP) averaged over occipito-parietal electrodes (P7, P3, P4, P8, POz, O1, O2, M1, and M2) show an early transient response followed by a sustained potential, which is stronger under high WM load (cluster permutation test, p < 0.05). **B**- Gaze velocity increases sharply following stimulus onset, peaking within the first second and remaining elevated for high compared to low WM load (cluster permutation test, p < 0.05). **C**- Head rotation velocity (Y-axis yaw) shows an initial rotation toward the stimulus followed by a return toward the road, with smaller corrective velocities for high WM load, indicating prolonged head orientation toward the stimulus (cluster permutation test, p < 0.05). **D**- Steering wheel deviation reflects greater modulation during high WM load (cluster permutation test, p < 0.05). **E**- Accelerator pedal pressure decreases more under high WM load, paralleling attentional engagement with the secondary task (cluster permutation test, p < 0.05). **F**- Vehicle speed shows a more pronounced decline for high WM load (cluster permutation test, p < 0.05). Shaded areas represent ±1 SEM across participants N=43.

Stimulus onset was further associated with systematic changes in driving behavior (Figure 2D-F). Steering wheel deviation (Figure 2D) showed a transient increase peaking around 2 s post-stimulus, followed by a compensatory shift in the opposite direction.There was no condition difference in terms of the size of this shift, except the delayed peak latencies for the high load condition. Similarly, accelerator pedal pressure (Figure 2E) decreased following stimulus onset, reaching its lowest point around 2-3 s. The reduction was larger for the high WM load condition, indicating a tendency for drivers to decelerate more when cognitive demands increased (cluster-based permutation test, *p* < 0.05). As a consequence of these adjustments, vehicle speed (Figure 2F) declined progressively throughout the trial. Speed reduction was more pronounced under high WM load, reflecting a compensatory strategy to maintain control during the cognitively demanding secondary task (cluster-based permutation test, *p* < 0.05). Overall, based on the peak latencies of gaze, head kinematics, and driving data, it becomes apparent that reaction peaks occur faster under low load compared to high load, consistent with the button-press reaction times observed later in the trial. These changes also prompted slight deviations in lateral distance from the midline, which were consistent across both conditions and remained within a mean range of approximately ±2 cm. Figure S2 in the Supplemental material provides further statistical details supporting these observations.

The time courses of both biological and driving data confirmed the effectiveness of the experimental manipulation. Higher working memory (WM) load distraction was associated with slower reaction times, reduced accuracy, and an overall decrease in driving speed, indicating a measurable impact on both cortical and behavioral measures.

Building on these behavioral and physiological effects, the cortical dynamics underlying these changes using a time-frequency analysis were examined next (Figure 3). This revealed a decrease in 20-30 Hz motor beta power and a suppression of alpha 8-12 Hz power over occipito-parietal regions contralateral to the attended stimulus location (Figure 3). Source reconstruction localized these effects to bilateral motor cortices and the contralateral visuo-parietal cortex, respectively, consistent with their known roles in movement preparation and spatial attention. In addition, a transient increase in evoked beta power (∼15 Hz) was observed, peaking within the first second after stimulus presentation. Source reconstruction indicated that this effect originated from the cerebellum ipsilateral to head rotation and stimulus attention, as well as the posterior parietal cortex (Figure 3), suggesting the involvement of these regions in rapid visuomotor coordination during distraction.

**Figure 3:**
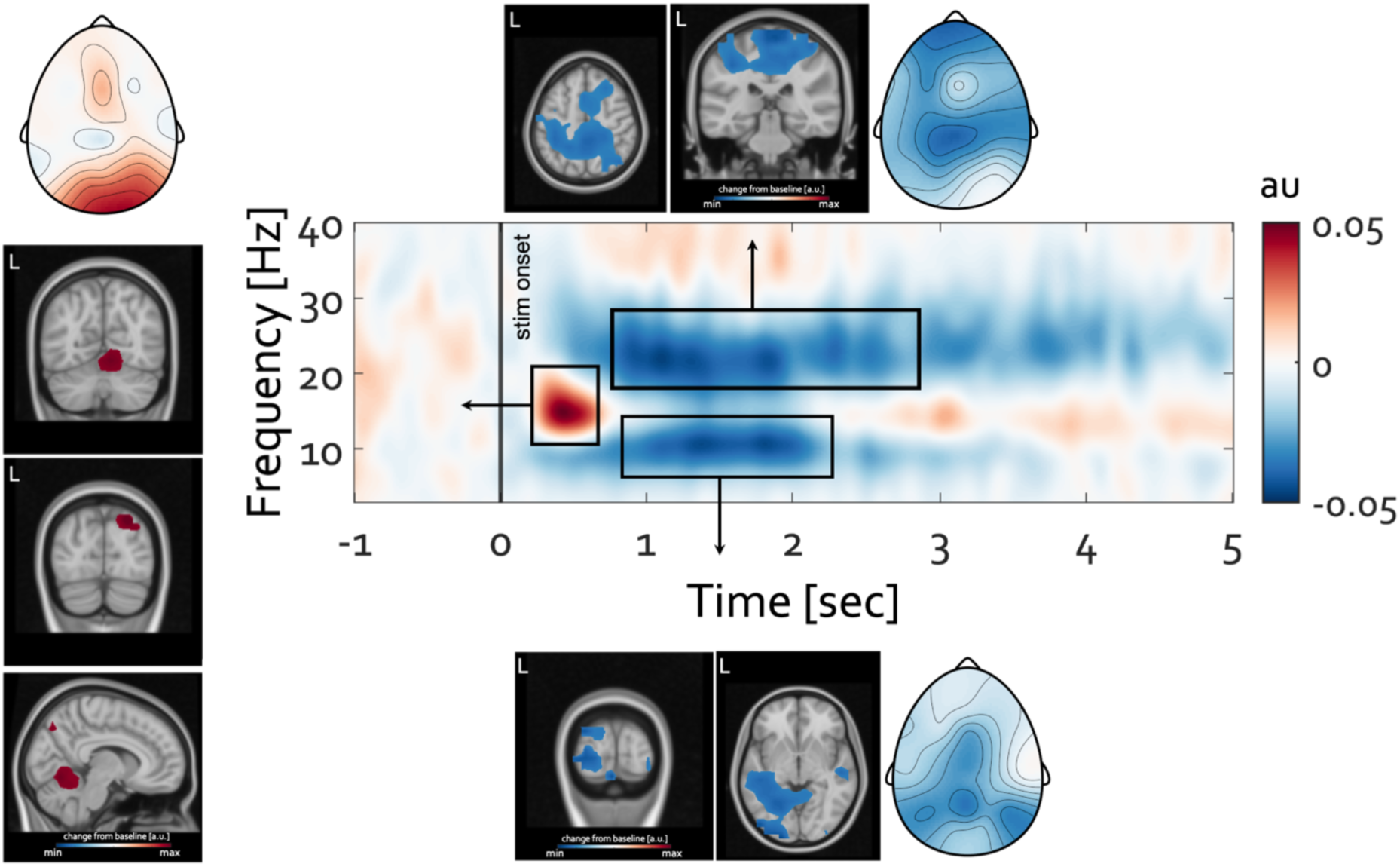
Source-resolved time-frequency dynamics of neural oscillations. Time-frequency representation over occipito-parietal electrodes(P7, P3, P4, P8, POz, O1, O2, M1, and M2), aligned to stimulus onset (0 s), shows power changes relative to a pre-stimulus baseline (warm colors: increases; cool colors: decreases). Black rectangles indicate the time-frequency windows illustrated in the scalp topographies and the source reconstruction maps. Source maps are thresholded at 80% of the respective maximum (for increases) or minimum (for decreases). Transient beta-band (∼15 Hz) power increases localized to the ipsilateral cerebellum and posterior parietal cortex (left), whereas sustained alpha (8-12 Hz) decreases originated from occipito-parietal cortex, and beta (15-25 Hz) decreases were observed over bilateral motor cortices.

After confirming that the experimental manipulation was effective and that the time-frequency data were reliably processed and source-imaged, the analysis focused on the main hypothesis. Based on previous findings linking alpha power modulation to attentional orienting and gaze control, it was predicted that alpha power would vary as a function of working memory load, with stronger alpha suppression expected for high-load stimuli, particularly in association with more sustained gaze shifts toward the stimulus.

These hypotheses are evaluated next. Figure 4 illustrates gaze position and time-frequency representations of oscillatory power changes for low and high working memory (WM) load conditions. During both loads gaze shifts were away from the road (blue) toward the stimulus position and return movements (red) are spatially consistent across trials with now statistically significant condition differences. For both conditions, stimulus onset was followed by a pronounced decrease in occipito-parietal alpha power (∼8-12 Hz) and a broader suppression of beta power (∼15-25 Hz). These effects appear stronger and more sustained for the high WM load condition compared to the low WM load condition, particularly in the alpha band, suggesting an increased allocation of visuospatial attention and greater task-related processing demands for high-load trials. More specifically, the alpha power modulation appears later in the course of the trial namely around 2 sec following stimulus onset.

**Figure 4:**
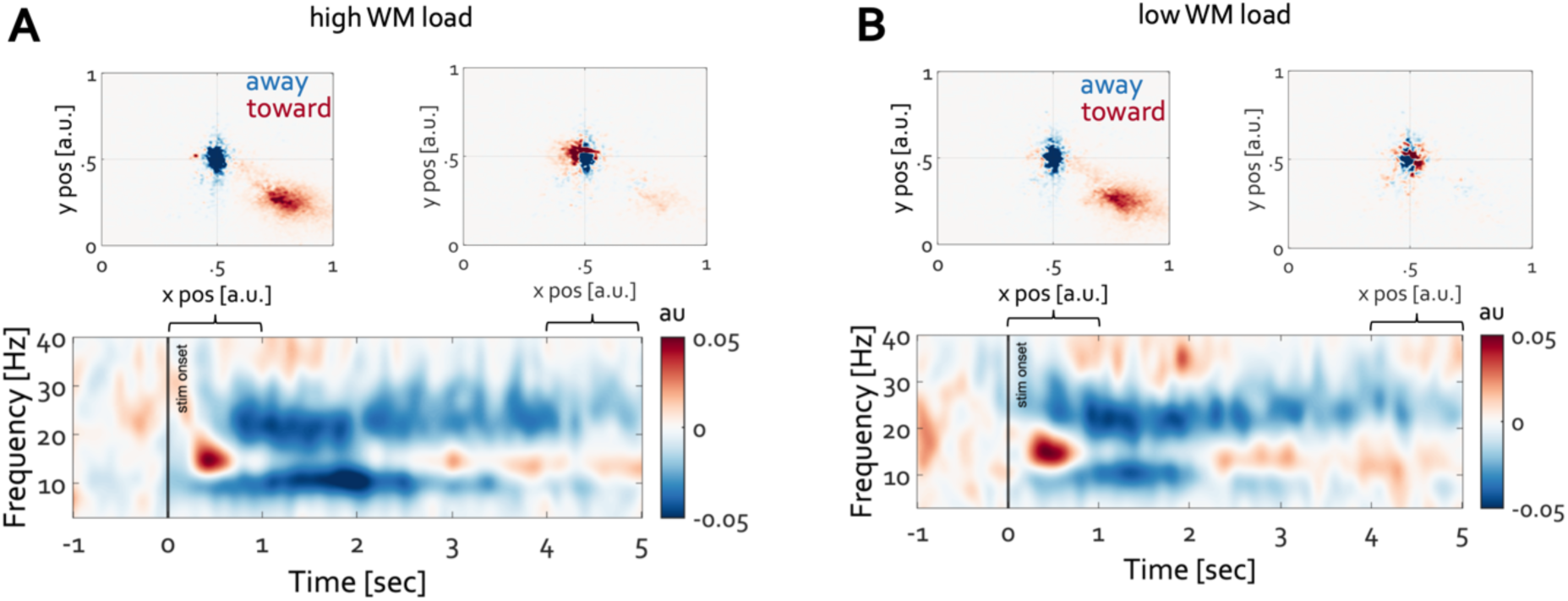
Gaze behavior and time-frequency dynamics as a function of working-memory (WM) load. **A**- high WM load. **B**- High WM load. Top panels show gaze distributions during the first (left) and last (right) second after stimulus onset, with red indicating gaze shifts toward the stimulus and blue indicating gaze shifts away from the road. Bottom panels display time-frequency representations of oscillatory power changes relative to a pre-stimulus baseline computed over electrodes P7, P3, P4, P8, POz, O1, O2, M1, and M2. Stimulus onset at 0 sec. Warm colors indicate power increases, and cool colors indicate decreases. Both conditions exhibit transient beta (∼15 Hz) increases followed by alpha (8-12 Hz) and beta (15-25 Hz) power decreases.

Indeed, both alpha power and gaze shifts exhibited statistically significant condition differences in this later time interval appr. 1.5 to 2.5sec after stimulus onset as illustrated in Figure 5A. The top row shows gaze distributions for high (left) and low (right) WM load conditions. Under high WM load, gaze remains directed longer and more consistently toward the stimulus location (red) as compared to low WM load. Cluster-based permutation test confirmed that the effect size of gaze differences between high and low WM load entails a cluster in the lower right visual field (Figure 5 middle, p < 0.05). This indicates that high WM load is associated with a stronger and more prolonged maintenance of gaze toward the stimulus location, in line with the time courses of the sustained ERP, gaze velocity and head rotation (Figure 2 A-C). The bottom panel of Figure 5 depicts the effect size of oscillatory power differences (high vs. low WM load) in the time-frequency domain. A robust cluster of alpha (8-13 Hz) power reduction is evident approximately 1.5-2.5 s after stimulus onset, with the scalp and source maps suggesting a distribution of this effect within occipito-parietal regions. Furthermore, as indicated by the sustained elevated levels of head movements toward the stimulus in the high-load condition (Figure 5B), the coupling between gaze shifts and parieto-occipital alpha power was preserved during the last second of the trial (Figure 5B, middle and bottom), albeit more weakly distributed within parietal cortical areas. To examine the temporal dynamics of gaze–alpha coupling, the gaze-density analysis was repeated using a 0.5-s sliding window centered on each time point. Horizontal gaze positions during fixations on the road and on the stimulus were extracted, transposed, and plotted as a function of time. Figure 6 shows the results. Raw power and power corrected for the aperiodic component over occipital–parietal electrodes are displayed for high- and low-load conditions alongside the time course of gaze shifts away from the road (Figure 6, middle) and toward the stimulus on the dashboard (Figure 6, bottom). Maintaining fixation on the road was associated with increased alpha power, whereas increased eye exploration was associated with decreased alpha power. The durations of these time courses closely matched the longer latencies observed for the high-load condition compared with the low-load condition (Figure 6, bottom).

**Figure 5:**
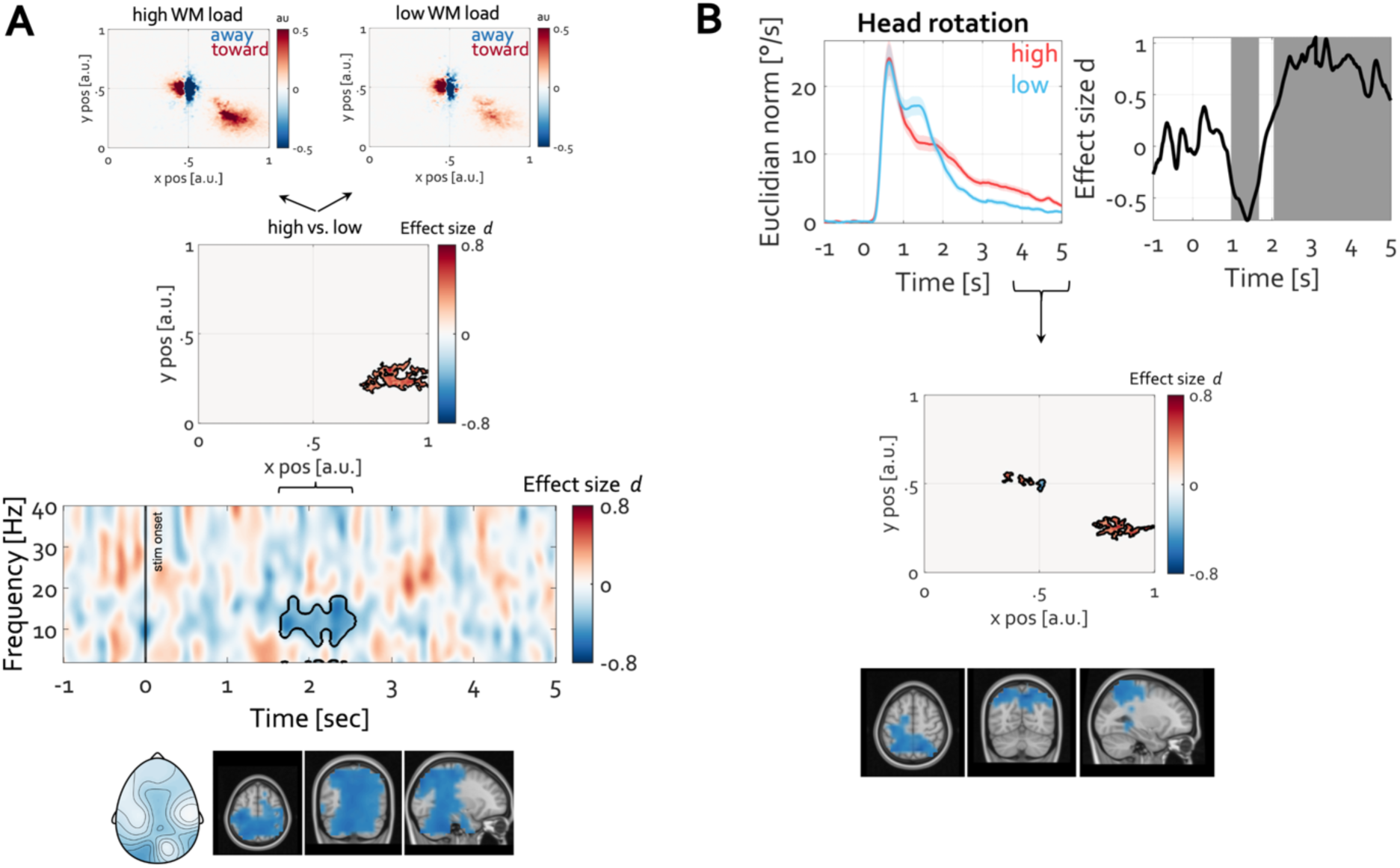
Load-dependent differences in gaze behavior and alpha power. **A-** Top: Gaze distributions within the time interval of 1.5 to 2.5 sec after stimulus onset for high (left) and low (right) WM load. Warm colors indicate gaze shifts toward the stimulus and cold colors indicating gaze shifts away. Middle: Effect size (Cohen’s *d*) map of gaze differences between high and low WM load, showing stronger gaze engagement toward the stimulus under high load. The black contour marks the cluster on the basis of which the null hypothesis of no condition difference was rejected (cluster-based permutation test, p = 0.002, clusterstat = 5.3223e+04). Bottom: Time-frequency representation of oscillatory power differences (high vs. low WM load) for electrode O1 expressed in units of Effect size d of the differences between high and low WM load. Warm colors represent higher effect size for high load, and cool colors indicate lower effect size for high load. The black contour marks the cluster on the basis of which the null hypothesis of no condition difference was rejected (cluster-based permutation test, p = 0.002, clusterstat = −105.2817). The scalp topography and source maps illustrate the distribution of this effect. Source maps thresholded at 5% alpha level (cluster-permutation test p= 0.0040, clusterstat = −2.6981e+03). **B-** Top: Time course of head rotation, incorporating all directions using the Euclidean norm (left), and the effect size of condition differences, with shaded areas denoting clusters supporting the conclusion of a significant effect. Bottom: As in A, but for the time interval 4-5 s within the trial.

**Figure 6:**
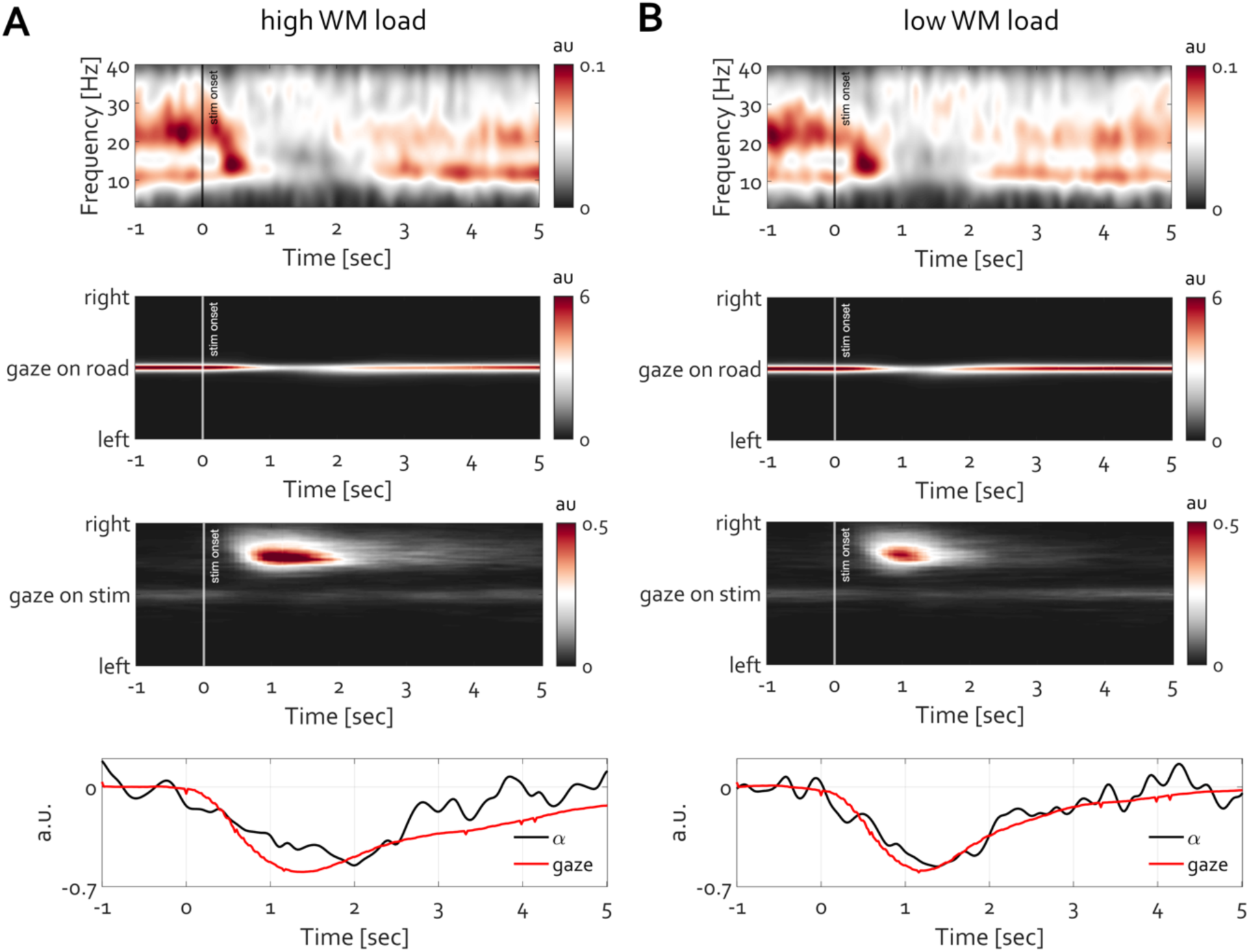
Time–frequency and temporally resolved gaze variability for high vs. low working memory (WM) load. **A-** High WM load and **B** low WM load conditions. Top row: grand averaged time-frequency representations (TFRs) of power (aperiodic component removed), aligned to stimulus onset (*t* = 0) and averaged across electrodes P7, P3, Pz, P4, P8, POz, O1, O2, M1, M2. Middle rows: horizontal gaze position variability over time for gaze on road and gaze on stimulus, with warmer colors indicating greater gaze concentration (arbitrary units, *au*). The temporal coincidence of reduced gaze on the road corresponds to increased gaze toward the stimulus located on the dashboard. Bottom row: average time courses of occipital alpha-band power (black) and gaze variability on the road (red), both aligned to stimulus onset, converted to normalized range-corrected values (−1 to 1) and expressed as change relative to the pre-stimulus baseline. Negative gaze values indicate a decrease in gaze maintenance on the road; as apparent from these illustrations, this decrease reflects gaze shifts toward the stimulus on the dashboard.

Finally, integrating all results provides a comprehensive view addressing the central research question: what are the neuro-ocular costs of texting while driving, and how do these costs manifest across perceptual, cognitive, and motor systems? Neurophysiological, ocular, and vehicle-control signals were aligned to stimulus onset (0 s) and baseline-normalized (−1 to 0 s) and are summarized in Figure 7A. In the high-load condition, eye velocity increased first, showing an initial peak at ∼200 ms. Perieto-cerebellar beta-band power (15-18 Hz) rose next, peaking at ∼400 ms, followed by a slow ERP component with a maximum at ∼980 ms. Around the same time, gaze shifted toward the stimulus on the dashboard. Changes in vehicle control occurred later: steering wheel deviation peaked at ∼1.84 s, and the mean button-press reaction time occurred at ∼4.5 s. Alpha power (8-12 Hz) was inversely related to exploratory gaze: periods of fixation on the road coincided with elevated alpha, whereas increased gaze exploration coincided with alpha decreases.

**Figure 7:**
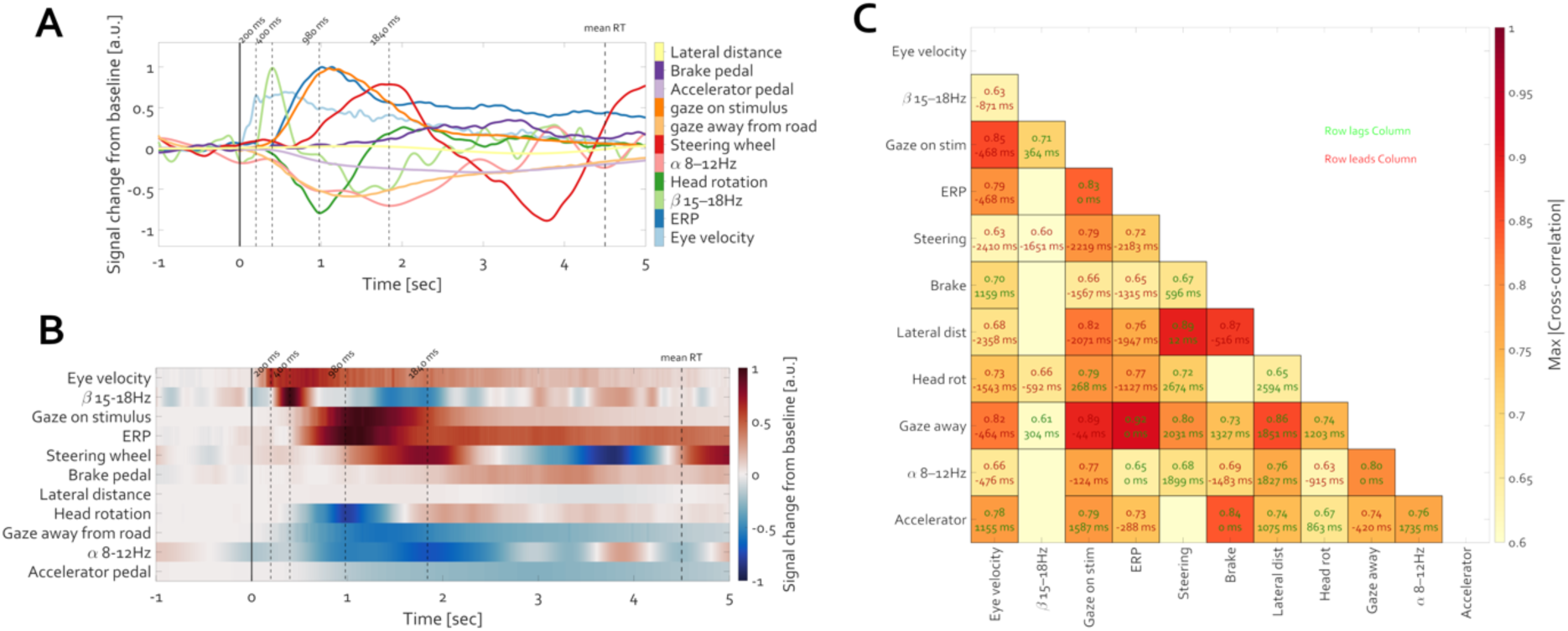
Neuro-ocular and behavioral dynamics during texting while driving. **A-**Normalized time courses of neurophysiological, ocular, and vehicle-control signals aligned to stimulus onset (0 sec). Illustrated is the high load condition. Vertical lines indicate key event-related response latencies: 200 ms first peak in eye velocity, 400 ms peak in beta-band (15-18 Hz) synchrony, 980 ms peak of slow ERP component and gaze shift to the stimulus on the dashboard, 1840 ms peak of steering wheel deviation and 4.5 sec - mean reaction time (RT) indicated by a button press. A broad set of signals are shown, including EEG activity (ERP, alpha, beta), gaze direction, head rotation, eye velocity, and driving behavior (steering, accelerator, brake, lateral distance). **B-** The same signals visualized as a heatmap to emphasize temporal dynamics and deviations from pre-stimulus baseline. Warm colors indicate increase (e.g. peaks in A) and cold colors a decrease (e.g. troughs in A) from pre stimulus baseline. **C-** Cross-correlation matrix showing peak absolute correlations between signals (color-coded) and the corresponding lag in milliseconds. Green labels indicate the row signal lags the column signal; red labels indicate the row signal leads the column signal. This matrix reveals the temporal coordination among perceptual, cognitive, and motor systems involved in driving and multitasking.

Pairwise cross-correlations (max |r| displayed) revealed strong coupling among modalities, with numerous pairs exceeding |r| ≈ 0.6–0.9. Lead–lag annotations (ms) indicated a sequential progression: ocular signals tended to precede cortical responses (beta, ERP), which in turn preceded head rotation and steering adjustments; downstream effects were observed in pedal activity and lateral distance. This pattern reflects coordinated processing across perceptual, cognitive, and motor systems during dual-task driving under high load.

In summary, these results show a consistent temporal cascade initiated by eye movements, followed by parieto-cerebellar beta and sustained ERP responses, and culminating in head and steering adjustments and broader vehicle-control changes. Alpha power tracked gaze strategy, increasing during road fixation and decreasing during exploratory gaze. Together, these findings delineate the timing and coupling of neuro-ocular and driving behaviors during texting while driving.

## Discussion

The present study demonstrates that, during a naturalistic simulated driving task, increasing WM load introduced via interaction with dashboard media systematically modulated behavioral, ocular, and neural measures. High WM load resulted in slower reaction times, reduced accuracy, prolonged gaze toward secondary stimuli, smaller corrective head rotations, and compensatory driving adjustments such as accelerator pedal mediated speed reduction. These behavioral changes were preceded by robust neural signatures, confirming that the experimental manipulation effectively induced cognitive distraction, even in a less artificially reduced, naturalistic setting, relevant for subsequently studying real-world risks of distraction in traffic.

### Parsimonious Interpretation of Alpha Modulation via Gaze Dynamics

Time-frequency analysis revealed a strong suppression of alpha power (∼8-13 Hz) over occipito-parietal regions contralateral to the stimulus, which scaled with WM load and co-occurred with sustained gaze shifts toward the stimulus. If access to eye-tracking and head-kinematics data had been absent, the present results could be interpreted entirely within dominant accounts of alpha oscillations. Specifically, the observed modulation of posterior alpha power with working-memory (WM) load would fit well with the “gating-by-inhibition” and “inhibition-timing” frameworks, whereby alpha increases serve to suppress processing in task-irrelevant regions, while alpha decreases disinhibit task-relevant areas, thereby facilitating information processing^6,7^. Such a view aligns with inhibition-timing accounts and more recent rhythmic gating formulations (e.g.,^15^), in which alpha provides a temporal control signal over the flow of sensory and mnemonic information. Within this framework, higher WM load would entail stronger suppression of irrelevant channels and more pronounced facilitation of relevant ones, leading to more efficient coding in the network that supports WM maintenance. However, recent perspectives^16^ caution against a single suppression-based interpretation. Foster and Awh (2019) emphasize that the existing evidence is equally compatible with an account in which alpha activity enables spatial and mnemonic selection through signal enhancement at relevant locations, rather than active suppression of irrelevant input. From this standpoint, posterior alpha modulations may track the locus and precision of covert orienting, without necessarily implying inhibitory control over unattended regions.

The complementary view afforded here by incorporating gaze and head-motion measures indicates that part of the alpha modulation can be attributed to overt and covert orienting behavior, as well as changes in visual input sampling across WM loads. This multimodal approach illustrates not only that alpha power reflects internal prioritization of information, but also that it is shaped by dynamic sensorimotor states that co-vary with cognitive demands. In doing so, it extends the interpretation beyond purely neural gating or suppression mechanisms, and towards a richer account in which alpha dynamics emerge from the interplay between cognitive control processes and embodied sampling of the environment. Specifically, the present results align with a growing body of evidence demonstrating that alpha oscillations are closely linked to oculomotor activity^17,23,34–37^ and gaze control rather than purely “top-down” cognitive inhibition. By incorporating concurrent gaze and head-tracking data, the present study provides a more parsimonious account of the robust alpha power suppressions observed under high working-memory load. Rather than invoking abstract “top-down” inhibitory control, the reductions in occipito-parietal alpha can be understood as a direct consequence of active visuomotor engagement - essentially reflecting neural regulation of the oculomotor system during increased task demand. This interpretation aligns with the perspective that the default mode of the human visual system is continuous eye movement (saccades, microsaccades, drifts) and that alpha oscillations serve to modulate this intrinsic activity. In the present driving paradigm, high cognitive load induced prolonged stimulus-directed gaze along with contralateral alpha power reduction, consistent with the idea that greater visual exploration (even if gaze is held on a screen, micro-movements and attentional shifts still occur) is associated with stronger alpha suppression. Notably, this sensorimotor framing circumvents the need for a homuncular “central executive” explanation of alpha changes. Instead, alpha power modulations are grounded in observable gaze behavior: overt action entails an increase in alpha power to stabilize vision when the degrees of freedom of gaze variation must be reduced in order to maintain relatively „fixed” position on the road ahead, and conversely, alpha power drops whenever exploratory eye movements and attentional shifts increase. The present findings thus reinforce an action-based mechanism for alpha oscillations in realistic tasks, echoing emerging models that situate alpha’s role in the active control of gaze rather than in disembodied cognitive inhibition^17,23^.

This integrated neuro-ocular approach fills critical gaps left by existing reviews of driver distraction and cognitive workload. For instance, Yusoff et al.^2^ emphasized the value of hybrid measures - combining eye-tracking indices with EEG - in detecting driver cognitive load, noting that such multimodal metrics are more sensitive and reliable than either modality alone. Likewise, Li et al.^41^ argued that interpretations of EEG rhythms during complex tasks must account for eye movement dynamics, as unrecognized oculomotor influences can confound cognitive inferences. Both reviews advocated for approaches that integrate gaze behavior with neural data to improve ecological validity and explanatory power. However, until now, empirical demonstrations of this principle in realistic settings have been limited. The present study directly answers these calls by showing that alpha oscillation changes and gaze behavior are tightly coupled during a real-life-mimicking driving task. In doing so, it extends the literature beyond prior lab-based findings and theoretical proposals. Present results empirically confirm that combining EEG with eye and head kinematics yields deeper insight into driver distraction: the close alpha-gaze correspondence reported here provides concrete mechanistic evidence addressing how and why cognitive load impacts the sensorimotor orchestration of attention in naturalistic context.

### Transient Cerebellar Activity and Gaze Control

A novel contribution of the present study is the identification of a cerebellar transient (∼15 Hz beta) oscillatory component linked to the gaze and head direction shifts under cognitive load. This finding was not anticipated by existing reviews of EEG in driving (which have focused mostly on cortical alpha/theta changes and behavioral metrics) and thus broadens the scope of known neural correlates of distraction. The transient beta increase localized in the vicinity of the ipsilateral vestibulo-cerebellum (flocculus/paraflocculus and oculomotor vermis) prior to each head rotation suggests the cerebellum’s involvement in preparing and refining gaze and head shifts. Unlike the cortex, which has both ipsilateral and contralateral connections, the cerebellum’s primary influence is on the side of the body that corresponds to the cerebellar hemisphere^42^. This is in line with longstanding neurophysiological evidence that the cerebellum aids gaze stabilization and eye-head coordination through predictive control mechanisms^43–45^. Indeed, the cerebellar signal observed here may reflect an anticipatory calibration of the vestibulo-ocular reflex and other oculomotor circuits, ensuring that diverting one’s gaze to an in-car display does not catastrophically impair spatial orientation and thus, constitutes a potentially previously unrecognized aspect of the driver’s neurophysiology. The implication is that attentional gaze shifts under load engage a distributed network that includes the cerebellum and posterior parietal circuits as a key node for sensorimotor integration. This insight connects with clinical and fundamental work on cerebellar contributions to attention and eye movement control^46–50^, suggesting that transient synchronous activity in this region could serve as an indicator for the timing of gaze-head adjustments. In summary, present results foreground the cerebellum’s role in the real-time guidance of action in service of visual attention-an advance in understanding that complements the cortical-focused discussions in earlier literature.

### Neuro-ocular cost of texting while driving

The present results reveal a clear temporal cascade of neural, ocular, and vehicular changes during high working memory (WM) load distraction in simulated driving. The sequence begins with a rapid rise in eye velocity, peaking at ∼200 ms after stimulus onset, marking the initiation of a gaze shift toward the stimulus on the dashboard. Around 400 ms, parieto-cerebellar beta-band power (15–18 Hz) peaks, followed by the emergence of a slow occipito-parietal ERP component, which reaches its maximum at ∼980 ms, coinciding with sustained gaze on the stimulus. Head rotation toward the stimulus peaks at ∼1 s, and steering deviation reaches its maximum at ∼1.84 s. Alpha power over occipito-parietal regions dynamically tracks gaze strategy - increasing during fixation on the road and decreasing during exploratory gaze toward the stimulus - with the high-load condition showing longer and more stable off-road gaze accompanied by sustained alpha suppression. High WM load also prolongs head orientation toward the stimulus and delays corrective head movement back to the road, as well as maintaining repeated small head adjustments later in the trial. Accelerator pedal pressure begins to decline within the first seconds after stimulus onset, reaching its lowest point between 2-3 s, with a more pronounced reduction under high load, leading to a progressive drop in vehicle speed. This event cascade occurs well before the mean button-press responses at ∼4.5 s. The cross-correlation analysis illustrated in Figure 7 confirms a lead-lag structure in which ocular changes precede cortical responses, which in turn precede head and steering adjustments, followed by pedal activity changes and lateral distance deviations.

Consequently, the cost of texting while driving extends well beyond the moment of visual diversion; degradation in performance begins before the gaze is fully shifted off-road and persists long after the eyes return to the roadway. Under high WM load, sustained off-road gaze is accompanied by prolonged alpha suppression, delayed corrective head and gaze movements, and extended periods of reduced visual sampling of the driving scene. Motor control adaptations - including reduced steering correction velocity, decreased accelerator pressure, and a progressive loss of speed - emerge downstream of these neural and ocular changes, reflecting a cascade of effects on vehicle handling. Early ocular shifts initiate a sequence of neural, head, and motor adjustments, creating a lag in overall driving readiness even after the secondary task has been completed.

These findings extend beyond traditional distraction metrics such as off-road glance duration by revealing a precise temporal cascade that links eye movements, cortical dynamics, head orientation, and vehicle control. The results demonstrate that alpha-band modulation closely tracks gaze strategy in realistic driving-like conditions, quantifying the sensory suppression associated with attention shifts away from the road. They provide direct evidence that high WM load prolongs both neural and behavioral costs of visual diversion, underscoring the interaction between cognitive load, visuomotor coordination, and driving performance. Importantly, they show that ocular and neural precursors to performance degradation arise before measurable changes in vehicle control, offering earlier indicators of distraction risk than those available through eye-tracking- only approaches.

### Insights of a multimodal naturalistic approach

By framing distraction in terms of coordinated neural and ocular dynamics, present study underscores the mechanistic insights enabled by multimodal analysis in naturalistic conditions. Traditional EEG studies both of cognitive load in laboratory settings and of driver workload have often interpreted alpha suppression as an isolated index of “mental effort”, but such interpretations can be ambiguous when divorced from behavior. Here, by concurrently monitoring where the eyes look and how the head moves, the presented results attribute meaning to alpha fluctuations in a concrete behavioral context. This represents both a methodological and conceptual advance. It demonstrates that what might superficially be labeled a purely “cognitive” EEG event is in fact deeply intertwined with action-specifically, the suspension or execution of gaze-head shifts. The present approach opens a more embodied account of cognitive load, that resonates with the default oculomotor mode of the human brain. Furthermore, the naturalistic driving scenario adds ecological validity that many prior lab studies lack. The ability to capture early neuro-ocular signatures of distraction before overt driving errors occur has practical implications for developing driver-monitoring systems. Overall, the present findings demonstrate the kind of rich, mechanistic understanding that can be gained by bridging neuroscience and behavior in complex settings. They extend and deepen prior work by concretely illustrating how multimodal data fusion leads to more parsimonious and actionable interpretations of driver distraction than either modality alone. By situating alpha suppression, gaze behavior, and cerebellar activity within the same experimental framework, the present study portrays a novel horizon for future research on action-attention coupling in real-world tasks, grounded firmly in an integrated neuroergonomic framework. Future work combining EEG, eye-tracking, and advanced source modeling may help establish these measures as robust biomarkers for attention-action coupling in applied and clinical settings.

### Limitations and Future Directions

Several limitations should be noted. First, although the study revealed robust coupling between alpha suppression and gaze behavior, these findings remain correlational. Future studies could employ gaze-contingent paradigms or brain stimulation to establish directionality. Second, the cerebellar beta activity identified through source reconstruction should be interpreted with caution. Although even a small number of electrodes (e.g. 19), when evenly distributed across the scalp, can yield similar distributions of source estimates compared to high-density setups^51^, complementary approaches such as MEG and high density EEG could provide stronger confirmation of this effect. Furthermore, although the driving simulator offered a controlled yet ecologically relevant environment, it cannot fully capture the complexity of real-world driving, including traffic dynamics and environmental unpredictability. Expanding the paradigm to naturalistic settings or advanced simulators would strengthen the ecological validity of the findings. In addition, the lack of direct manipulation of gaze behavior, the moderate sample size, and the absence of additional behavioral safety metrics (e.g., hazard response) limit the generalizability of the results.

Future research should therefore test whether alpha-gaze coupling and cerebellar beta activity generalize across different driving contexts, include causal manipulations, and examine whether these neural markers can predict real-world driving risk in more complex driving tasks. Such efforts may ultimately inform the development of neurophysiological monitoring tools for distraction detection and enhance our understanding of visuomotor attention in both applied and clinical contexts.

## Acknowledgments

This work was supported by the Schweizerischer Nationalfonds zur Förderung der Wissenschaftlichen Forschung (SNF) Grant 105314_207580 awarded to PT and Emma-Louise-Kessler Fund awarded to LS. The authors extend gratitude to all the participants who volunteered for the experiment.

## Methods

### Participants

A total of 43 participants were recruited. Participants were recruited from the local university and through community advertisements. The sample comprised 27 female and 16 male participants, with an age range of 22 to 55 years (M = 29 years). Prior to participation, all participants provided written informed consent in accordance with the Declaration of Helsinki. The studies were approved by the local ethics committee at the host institution-IRB number 25.05.15. Participants received monetary compensation for their participation. This study was not preregistered. The sample size was limited by resource constraints, such as time constraints as well as budget limitations. As argued by Lakens^52^, these potential limitations are acknowledged here, given the practical constraints preventing the achievement of an ideal sample size based on power analysis.

### EEG and Eye Tracking

Stimuli were displayed on a Fujitsu CELSIUS H Series monitor with 1920×1080 pixel resolution. Participants were seated in front of the steering wheel in a BMW i3 car. Stimuli were presented using PsychoPy ^53^ on a full-screen black background.

EEG data were acquired using the mBrainTrain Smarting device, a wireless 24-channel EEG system. The system utilizes semi-dry/saline-based Ag/AgCl electrodes embedded in a cap, adhering to the International 10-20 System of electrode placement. Electrodes were referenced online to the FCz electrode, with AFz serving as the ground. Impedance for each electrode was maintained below 10 kΩ. Data were acquired at 250Hz sampling rate and converted offline to an average reference for further analysis. In addition, data from an in-ear EEG device (IDUN Technologies) were collected from 27 participants, which is outside the scope of the present report and will not be addressed further.

Eye-tracking data were acquired using a Tobii Pro Glasses 3 eye tracker, with real-time gaze data transmitted via the Lab Streaming Layer (LSL) protocol. The eye tracker captured left and right gaze points and pupil diameter, in addition to gyroscope and accelerometer data. Triggers marking stimulus onset were sent via LSL using a separate stream for synchronization of the EEG and eye tracking and SILAB (see below) streams.

### Driving simulator

The VICTOR (Vehicle for Interdisciplinary Clinical and Translational Research) driving simulator is built around a complete BMW i3 chassis. Its original shock absorbers were replaced with custom mounts for spindle actuators (Festo, Esslingen, Germany; actuator ESBF-BS-50-100-20P, servomotor EMMB-AS-80-07-S30S, servo drive CMMT-AS-C4-3A-PN-S1; Neugart, Kippenheim, Germany, gear Art-No. 58474). These modifications, which include several specially manufactured components, enable the simulator to deliver motion feedback primarily for surface irregularities, acceleration, and braking. Standard mirrors and dashboard displays were substituted or concealed with LCD monitors (Beetronics 10HD7 or 13HD7). A 40-inch screen (IIYAMA X4071UHSU-B1) was installed in the trunk area to replicate the rear-view mirror display. Additionally, the center console features a touch display (Asus Zen Touch MB16AMT) for presenting and collecting supplementary information. The throttle, brake, and accelerator pedals, as well as the steering and indicator controls, were adapted to integrate seamlessly with the virtual environment (SILAB 7.0, WIVW, Würzburg, Germany). SILAB is responsible for vehicle dynamics simulation and rendering the virtual environment on the car’s internal displays and through five DLP Laser Phosphor projectors (Barco F-80_Q7 with GLD 0.85-1.06:1 optics). These projectors are mounted on custom position-control systems (Lumachroma, Switzerland) and project onto a 275° fixed drywall projection screen. The five WQXGA images (2,560 × 1,600) are processed using VIOSO AnyBlend (Vioso, Düsseldorf, Germany) for mapping, warping, and blending. Visual rendering and driving parameter recording are performed at 60 Hz. The system runs on ten identical computers (Intel Core i7-9700 @ 3.6 GHz, 16 GB RAM, NVIDIA GeForce RTX 2080 SUPER), all networked and controlled via SILAB from a single LAN-connected workstation (“kiosk”). The simulator room is climate-controlled at 22°C (range 20-24°C), and the car interior is ventilated with fresh air at 21°C supplied by the building’s main HVAC system. Most technical components are either hidden in the car’s trunk or placed out of sight, creating a minimalistic environment that resembles a stationary car in a garage, thereby lowering the psychological entry barrier for users. A virtual 360° tour of the setup is available online: https://my.matterport.com/show/?m=ntULEgy28cU.

### Driving scenario

A customized monotonous nighttime driving scenario was employed. It is based on ten modules derived from a validated vigilance-driving scenario included in the Swiss-adapted scenario package for driver fitness evaluation (SPCH-DFA, WIVW, Würzburg, Germany). The scenario consists of approximately 45 minutes of low-stimulation driving on a single-lane highway, where the participant follows a lead vehicle at a safe distance without overtaking. The lead vehicle alternates its speed between 80 and 90 km/h, while the posted speed limit remains 100 km/h. Oncoming vehicles appear every 5 to 20 seconds. The simulator’s speedometer is disabled to reduce light emissions and to encourage focus on the road. The motion platform provides feedback for road surface changes and speed variations, but no additional motion cues are generated for lane departures or collisions. Each of the ten scenario modules is individually assessed for driving quality using SAFE (Standardized Application for Fitness to Drive Evaluations), software directly linked to SILAB.

Driving metrics such as speed, headway distance, and standard deviation of lateral position (SDLP) are recorded in SILAB and streamed through LSL for post hoc analysis and synchronization with EEG and eye-tracking data. Furthermore, standard driving errors-including lane departures, near collisions, and collisions—are automatically logged by SAFE.

### Procedure

Participants were initially introduced to the simulator environment and given a brief 10-minute familiarization drive. This session was intended to ensure they were comfortable with the controls and to screen for potential simulator sickness. No participant reported any symptoms during this phase, likely due to the minimal and highly controlled nature of the driving scenario. Following familiarization, the experimental procedure was explained in detail. Participants were then provided with an opportunity to ask questions before signing the informed consent form. Once consent was obtained, the EEG cap was mounted, and electrode impedance was checked to ensure signal quality. The eye-tracking system (Tobii Pro Glasses 3) was subsequently fitted and calibrated to each participant’s gaze using the integrated calibration routine. After setup, participants were seated in the simulator, and a final synchronization check between EEG, eye tracking, and driving data streams (via Lab Streaming Layer) was performed. The experimental session then began with the monotonous nighttime driving task, during which arithmetic stimuli were presented in random order on a dashboard-mounted screen under low- and high-working-memory load conditions. Participants completed the driving task while EEG, eye tracking, head kinematics, and driving performance data were recorded continuously. Upon completion of the approximately 45-minute driving session, participants were debriefed, the equipment was removed, and they received monetary compensation of 40 CHF.

### Data Processing and Analysis

EEG and eye-tracking data were preprocessed offline using the open-source FieldTrip ^54^ toolbox for neurophysiological data analysis. Continuous recordings were segmented into epochs time-locked to stimulus onset, including a 2-second pre-stimulus baseline. A finite impulse response (FIR) band-pass filter (1-45 Hz) was first applied to the raw EEG data, followed by artifact correction using independent component analysis (ICA) to remove components associated with ocular and cardiac activity. The cleaned ICA components were then projected back onto the raw data. After artifact correction, a 45 Hz low-pass filter was applied to minimize line noise while retaining low-frequency neural signals.

Time-frequency power spectra were estimated using multitaper convolution with a Hanning taper and a fixed 0.5-second sliding window, yielding a frequency resolution of 2 Hz in the range of 2-40 Hz. The window was advanced in 50 ms steps across each epoch, covering −2 to 5 seconds relative to stimulus onset. Power values were averaged across trials for each condition and baseline-corrected using the interval from −∞ to −0.25 seconds before stimulus onset. To separate oscillatory from aperiodic activity, the 1/f component was removed on a trial-by-trial basis using the specparam algorithm, which parameterizes neural power spectra into their periodic and aperiodic components ^55^.

Source reconstruction was performed using the linearly constrained minimum variance (LCMV) beamformer ^56^. Spatial filters were derived from the data covariance matrix and forward models based on the MNI ICBM 2009 template brain with a three-layer boundary element model (BEM) and standard electrode positions. These filters were applied to obtain virtual source time series at 2461 equally spaced grid points. The resulting source-level signals were subjected to the same time-frequency analysis as the sensor-level data.

### Eye tracking Data Processing

Raw eye-tracking data were analyzed to compute three-dimensional gaze density maps and instantaneous gaze velocity. Gaze positions in horizontal (x) and vertical (y) coordinates were extracted for each time point and binned into a 1,000 × 1,000 grid. Using MATLAB’s histcounts2, 2D histograms of gaze density were generated and smoothed with a Gaussian filter (imgaussfilt, smoothing factor 5) to create continuous heat maps. These maps were converted into FieldTrip-compatible structures with smoothed gaze density stored in powspctrm, analogous to time-frequency EEG data, and horizontal and vertical positions represented as time and frequency axes, respectively. This format allowed statistical comparisons between gaze behavior and neural activity within the same analytical framework. Gaze velocity was derived from the temporal derivatives of horizontal and vertical gaze positions. The signals were smoothed using a 100 ms moving average window (25 samples) before differentiation. Absolute velocity values were then stored in FieldTrip format, enabling direct integration of gaze velocity, gaze density, and EEG data for statistical analysis across experimental conditions.

### Statistical Analysis

To control for multiple comparisons, statistical testing was performed using a cluster-based permutation approach ^57^. This method accounts for spatial, temporal, and frequency correlations by identifying clusters of adjacent significant data points. A two-tailed alpha threshold of 0.05 and 1000 permutations were used to assess statistical significance.

### Data and Code Availability

All data and the code necessary to reproduce the present results will be made available upon publication.

