## Supplemental material for "The neuro-ocular costs of texting during driving"

**Supplementary material**

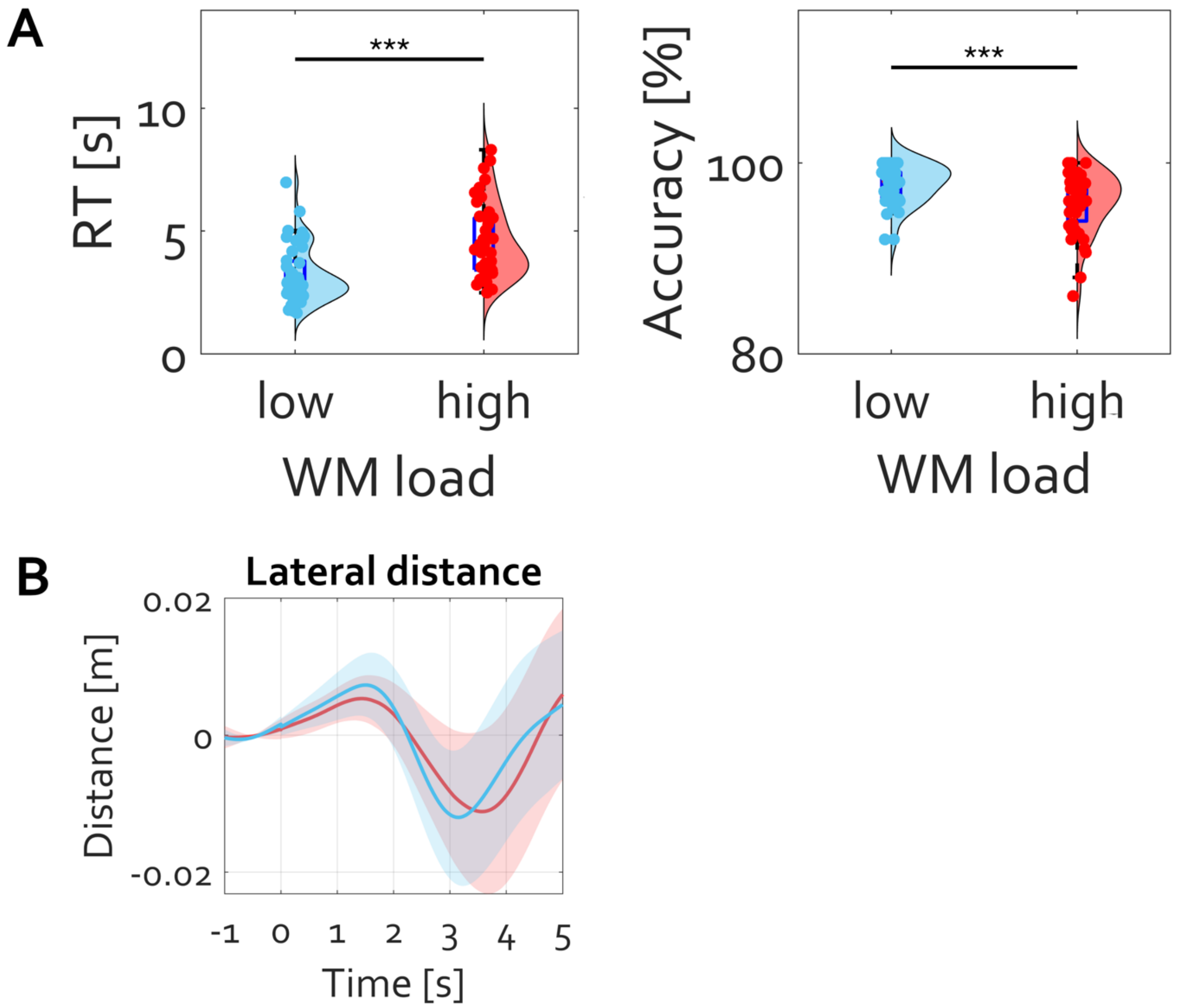


**Figure S 1: Effects of working memory (WM) load on behavioral performance and lateral vehicle control.**
(A) **Reaction time (RT) and accuracy:** Violin plots show RT (left) and accuracy (right) for low and high WM load conditions. Participants responded significantly slower and less accurately in the high WM load condition compared to the low WM load condition ($p < 0.001$).
(B) **Lateral distance:** Time-resolved lateral distance relative to the lane center for low (blue) and high (red) WM load conditions. Shaded areas represent the standard error of the mean (SEM).

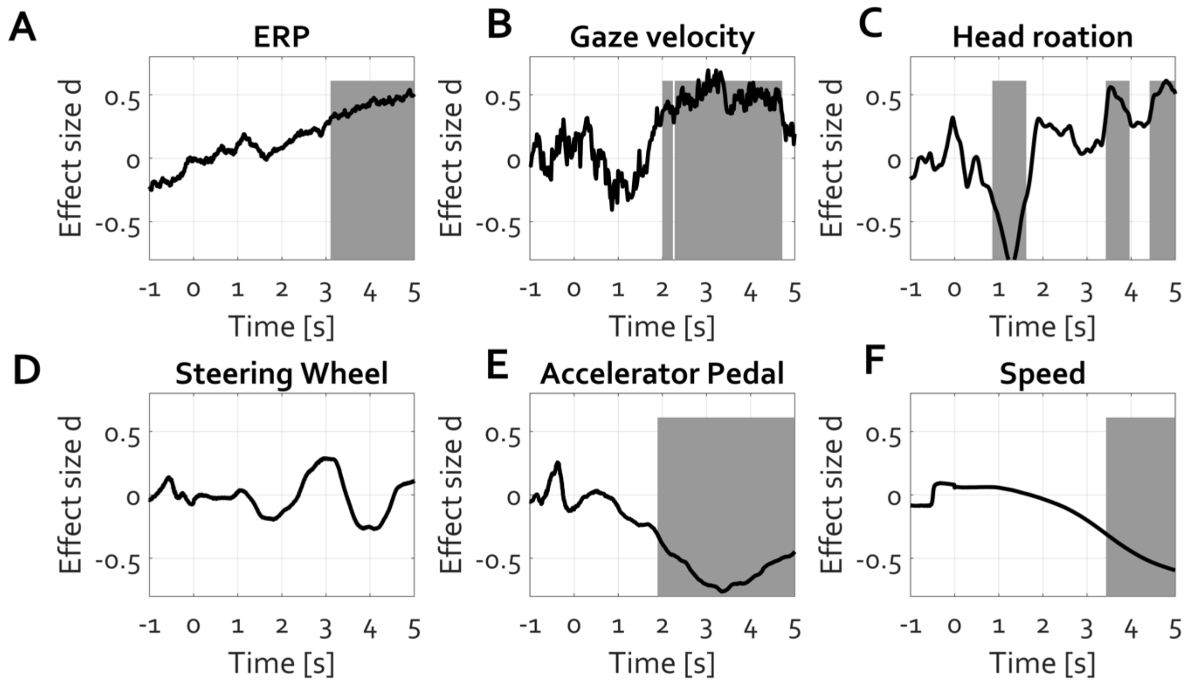


**Figure S 2: Time-resolved effect sizes (Cohen's *d*) for different behavioral and physiological measures.**
(A) **ERP:** Event-related potentials show a positive cluster from ~3 to 5 s (gray shaded area; $p = 0.006$, clusterstat = $1.304 \times 10^3$, stddev = 0.0035, cirange = 0.0068) based on which the null hypothesis ($H_0$) of no condition differences was rejected. (B) **Gaze velocity:** A positive cluster from ~2.3 to 5 s ($p = 0.002$, clusterstat = 393.99, stddev = 0.002, cirange = 0.0039) indicates time periods where $H_0$ was rejected.
(C) **Head rotation:** One negative cluster around 1–2 s ($p = 0.002$, clusterstat = –350.93, stddev = 0.002, cirange = 0.0039) and two positive clusters around 1–2 s ($p = 0.006$, clusterstat = 237.55, stddev = 0.0035, cirange = 0.0068) and 3.5–5 s ($p = 0.010$, clusterstat = 191.48, stddev = 0.0044, cirange = 0.0087) were identified, each rejecting $H_0$. (D) **Steering wheel:** No clusters where $H_0$ was rejected were observed.
(E) **Accelerator pedal:** A negative cluster from ~2.5 to 4.5 s ($p = 0.002$, clusterstat = $-1.461 \times 10^3$, stddev = 0.002, cirange = 0.0039) was identified where $H_0$ was rejected.
(F) **Speed:** A negative cluster from ~3.8 to 5 s ($p = 0.0299$, clusterstat = –588.51, stddev = 0.0076, cirange = 0.0149) indicates time periods where $H_0$ was rejected.
